# Systems Analysis of Subjects Acutely Infected with Chikungunya Virus

**DOI:** 10.1101/531921

**Authors:** Alessandra Soares-Schanoski, Natália Baptista Cruz, Luíza Antunes de Castro-Jorge, Renan Villanova Homem de Carvalho, Cliomar Alves dos Santos, Nancy da Rós, Úrsula Oliveira, Danuza Duarte Costa, Cecília Luíza Simões dos Santos, Marielton dos Passos Cunha, Maria Leonor Sarno Oliveira, Juliana Cardoso Alves, Regina Adalva de Lucena Couto Océa, Danielle Rodrigues Ribeiro, André Nicolau Aquime Gonçalves, Patricia Gonzalez, Andreas Suhrbier, Paolo Marinho de Andrade Zanotto, Inácio Junqueira de Azevedo, Dario S. Zamboni, Roque Pacheco Almeida, Paulo Lee Ho, Jorge Kalil, Milton Yutaka Nishiyama Junior, Helder I Nakaya

**Affiliations:** Bacteriology Laboratory, Butantan Institute, Brazil; Department of Clinical and Toxicological Analyses, School of Pharmaceutical Sciences, University of São Paulo, São Paulo, Brazil; Departamento de Biologia Celular, Molecular e Bioagentes Patogênicos, Faculdade de Medicina de Ribeirão Preto, Universidade de São Paulo, Ribeirão Preto, Brazil; Health Foundation Parreiras Horta (FSPH), Central Laboratory of Public Health (LACEN/SE), State Secretary for Health, Brazil; Special Laboratory for Applied Toxinology, Butantan Institute, Brazil; Respiratory Diseases Division, Virology Center, Adolfo Lutz Institute, Sao Paulo, SP, Brazil; Laboratory of Molecular Evolution and Bioinformatics, Department of Microbiology, Biomedical Sciences Institute, University of São Paulo, São Paulo, Brazil; Division of Immunology and Molecular Biology Laboratory, University Hospital/EBSERH, Federal University of Sergipe (UFS), Brazil; QIMR Berghofer Medical Research Institute, Brisbane, Queensland, Australia; Bacteriology Service, Bioindustrial Division, Butantan Institute, Brazil; Heart Institute, Faculty of Medicine, University of São Paulo (USP), Brazil

**Keywords:** Chikungunya, Systems Biology, Transcriptome

## Abstract

The largest ever recorded epidemic of the chikungunya virus (CHIKV) began in 2004 and affected four continents. Acute symptomatic infections are typically associated with the onset of fever and often debilitating polyarthralgia/polyarthritis. In this study, a systems biology approach was used to analyze the blood transcriptomes of adults acutely infected with CHIKV. Gene signatures that were associated with viral RNA amounts and to the onset of symptoms were identified. Among those genes, the putative role of Eukaryotic Initiation Factor (eIF) family genes and apolipoprotein B mRNA editing catalytic polypeptide-like (APOBEC3A) in the CHIKV replication process were displayed. We further compared these signatures with those induced by dengue virus infection and rheumatoid arthritis. Finally, we demonstrated that CHIKV infection in mice induced IL-1 beta production in a mechanism highly dependent on the inflammasome NLRP3 activation. The findings provided valuable insights into the virus–host interactions during the acute phase and could be useful in the investigation of new and effective therapeutic interventions.

## Introduction

Chikungunya virus (CHIKV) is a mosquito-borne reemerging arbovirus responsible for intermittent and devastating outbreaks (1). The largest epidemic of CHIKV ever recorded started in Africa in 2004 and has spread globally, reaching the Americas in 2014. The disease has reached four continents, affected more than 100 countries and infected over 10 million people (2, 3). CHIKV has spread rapidly through several Brazilian states and resulted in a total of 20,598 infected individuals in 2015 (4) and more than 200,000 suspected cases reported in 2017–2018 (5).

CHIKV acute infection typically results in a 5–7 days viremia and the symptoms are characterized by fever, rash and severe polyarthralgia/polyarthritis that can become chronic and persist from months to years (6). Mortality rates are estimated to be approximately 0.1% (7); however, the high attack rates (up to 30–75%) can result in a considerable economic burden (8, 9). The current mainstay of treatment involves the use of NSIADs and/or acetaminophen, although relief is often inadequate and more effective treatments are actively being sought (10). There has been a considerable quest to better understand CHIKV disease pathogenesis and virus–host interactions (11) using *in vitro* approaches (12) or animal models (13, 14).

Systems biology approaches have been successfully applied to identify molecular signatures associated with infections (15) and vaccination (16). In this study, a systems approach was performed to investigate subjects naturally infected with CHIKV during the 2016 epidemics in the state of Sergipe, Brazil. The blood transcriptome analyses performed in this study revealed key genes and pathways involved in acute CHIKV infection, providing important insights into how CHIKV interacts with the host’s immune system. These analyses helped uncover potential drug targets for improving CHIKV therapy.

## Results

### Clinical information and diagnosis of subjects infected with CHIKV

In an endemic area for CHIKV, Zika and dengue virus (17) in the northeast of Brazil, whole blood samples were collected from 39 adults with symptoms consistent with arboviral infection including fever, arthralgia, headache and/or muscle pain. Most individuals (n = 22) reported that the onset of the symptoms occurred on the same day or one day prior to the blood collection, indicating a very early stage of the disease (Fig. 1a and Table S1). Another eight had symptoms for two days and seven of them showed symptoms between three and four days. Two of the subjects reported pain for 20 days or more (Fig. 1a). Blood was also collected from 20 healthy subjects with no symptoms of viral infection and who tested negative for CHIKV RNA from endemic and non-endemic regions in Brazil. Serodiagnostic ELISAs were also performed to detect the presence of IgG antibodies specific to Zika and/or dengue viruses (Fig. 1b). We detected IgG and IgM antibodies specific to CHIKV in only one subject. The presence of CHIKV RNA by real-time RT-PCR or IgM for CHIKV positivity data, and the absence of Zika and dengue RNA were confirmed in the serum of all the patients (Table S1). The most frequent symptoms of CHIKV-infected patients were fever, arthralgia, headache and myalgia (Fig. 1c). As expected, individuals who reported being in the very early stage of the disease (Fig. 1d) had the highest levels of serum CHIKV RNA.

**Figure 1.**
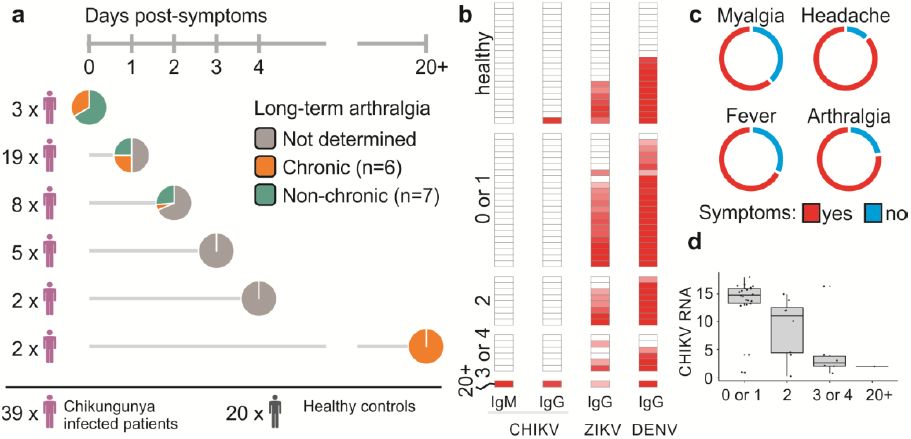
CHIKV samples collection, clinical data and diagnostics. (a) Study design representing the number of individuals and the days after the onset of symptoms that the blood samples were collected. The status of the long-term arthralgia chronicity was assessed in some of the patients as indicated by the following color scheme: chronic (orange), non-chronic (green) and undetermined (gray). (b) Serological diagnostics for CHIKV, DENV and ZIKV showing highest patients’ immunity for dengue and Zika in red scale (n=37). (c) Doughnut chart indicating the most prevalent symptoms reported by the patients. (d) CHIKV RNA (inverted Ct) in the subgroups determined by the day after the onset of symptom that the samples were collected (n=37).

### The impact of CHIKV acute infection on peripheral blood transcriptome

To understand the molecular changes that occurred during acute CHIKV infection, we compared the blood transcriptomes of infected individuals with those from healthy controls. Although the CHIKV RNA amounts had a small contribution to the variance, individuals were not grouped by the period of the onset of symptoms nor their infection status (Fig. S1b).

**Figure S1.**
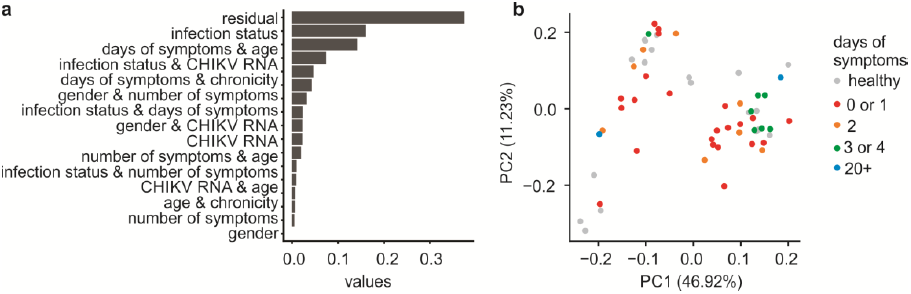
Unsupervised analysis highlights that the variation among individuals can be explained by undefined residual effects. (a) Histogram representing the effects ordered according to their contribution to the sample’s variance measured by PVCA. (b) Unsupervised principal components analysis (PCA) of the 59 patients, classified according to the expression data of healthy subjects and CHIKV-infected samples according to their days of symptoms represented by the two-dimensional PCA.

Correlation analysis between gene expression and the amounts of CHIKV RNA in the peripheral blood of infected individuals revealed approximately 3,500 genes associated with CHIKV infection. The expression of most of these genes (> 85%) was negatively correlated with CHIKV RNA (Fig. 2a). Using the correlation values as rank and Blood Transcription Modules (BTM) as gene sets, we run a gene set enrichment analysis (GSEA) to reveal the BTMs related to the present amount of CHIKV virus. BTMs associated with innate immune cells (dendritic cells and neutrophils), antiviral response, inflammation and toll-like receptor signaling (Fig. 2b and S4) were positively associated with the amount of CHIKV RNA. BTMs negatively associated with the amount of CHIKV RNA, i.e. with a negative normalized enrichment score (NES) were related to adaptive immune response, such as cell proliferation, activation and differentiation of B and T lymphocytes, in addition to chemokines and cytokines that could be due to the CHIKV infection related lymphopenia (Fig. 2b) (18). A gene network was subsequently constructed by connecting the genes related to BTMs with negative NES score (Fig. 2b) utilizing protein-protein interaction (PPI) data from InnateDB. Genes related to adaptive immune response and cells recruitment, such as *CCL5, CD8A* and *CD8B, CD3D, CD19, CCR7 and TCF7* among others appear in the respective network (Fig. 2c). Several of these genes are related to the homing/recruiting of the effector/memory T lymphocytes as well as to activation and induction of memory cells. The same process was performed for BTMs with positive NES score (Fig. S2). Genes such as *IRF7, OAS3, IFIT2* and *IFIT3, CXCL11, SERPING1* are all related to the innate immune response to viral infections (19) or can even be considered as markers of viral infections and some are ISGs (interferon-stimulated genes).

**Figure 2.**
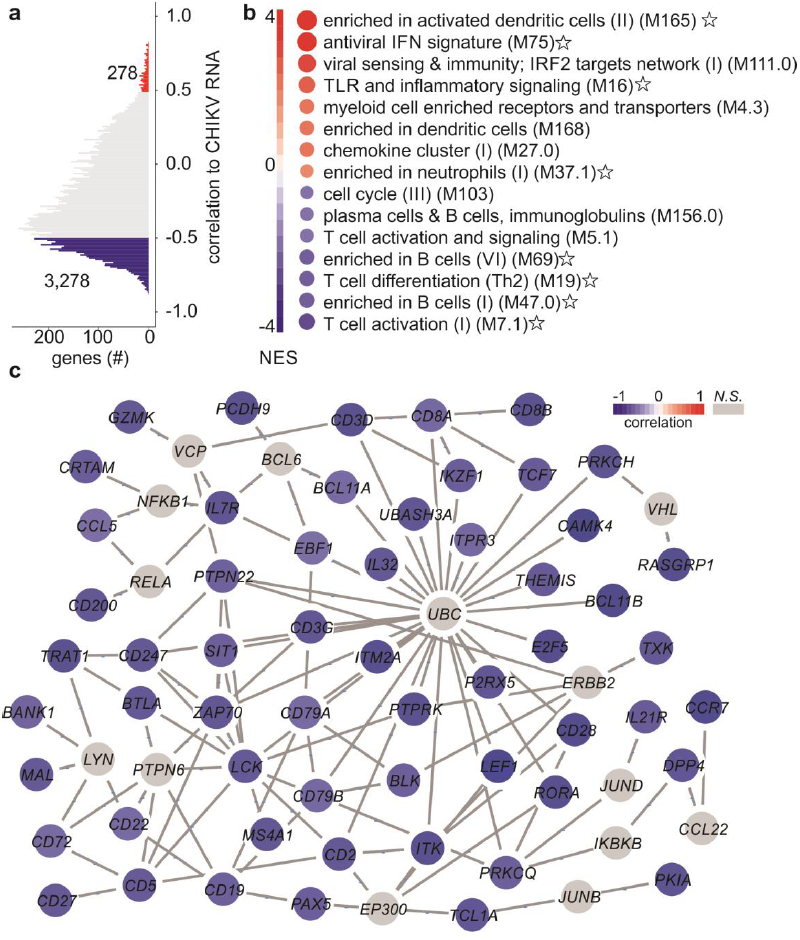
Gene signatures associated with the amounts of CHIKV RNA. (a) Representation of genes whose expression were correlated (Pearson’s r) with the amounts of CHIKV RNA (inverted Ct) and the respective number of genes with p-adjusted value < 0.01 and |R| > 0.5. (b) GSEA using the correlation values between expression and inverted Ct as ranks and BTMs as gene sets. The stars correspond to the gene sets used to construct the networks shown in panel C (negatively correlated pathways) and Fig. S2 (positively correlated pathways). (c) Minimum network constructed using the gene sets that presented a negative NES score with NetworkAnalyst tool. The protein-protein interaction (PPI) data was based on InnateDB and the graph was generated with Cytoscape tool. Blue nodes represent genes negatively correlated with amounts of CHIKV RNA and the color scheme indicates its strength. The gray nodes were added by NetworkAnalyst and are not part of the correlated genes.

**Figure S2.**
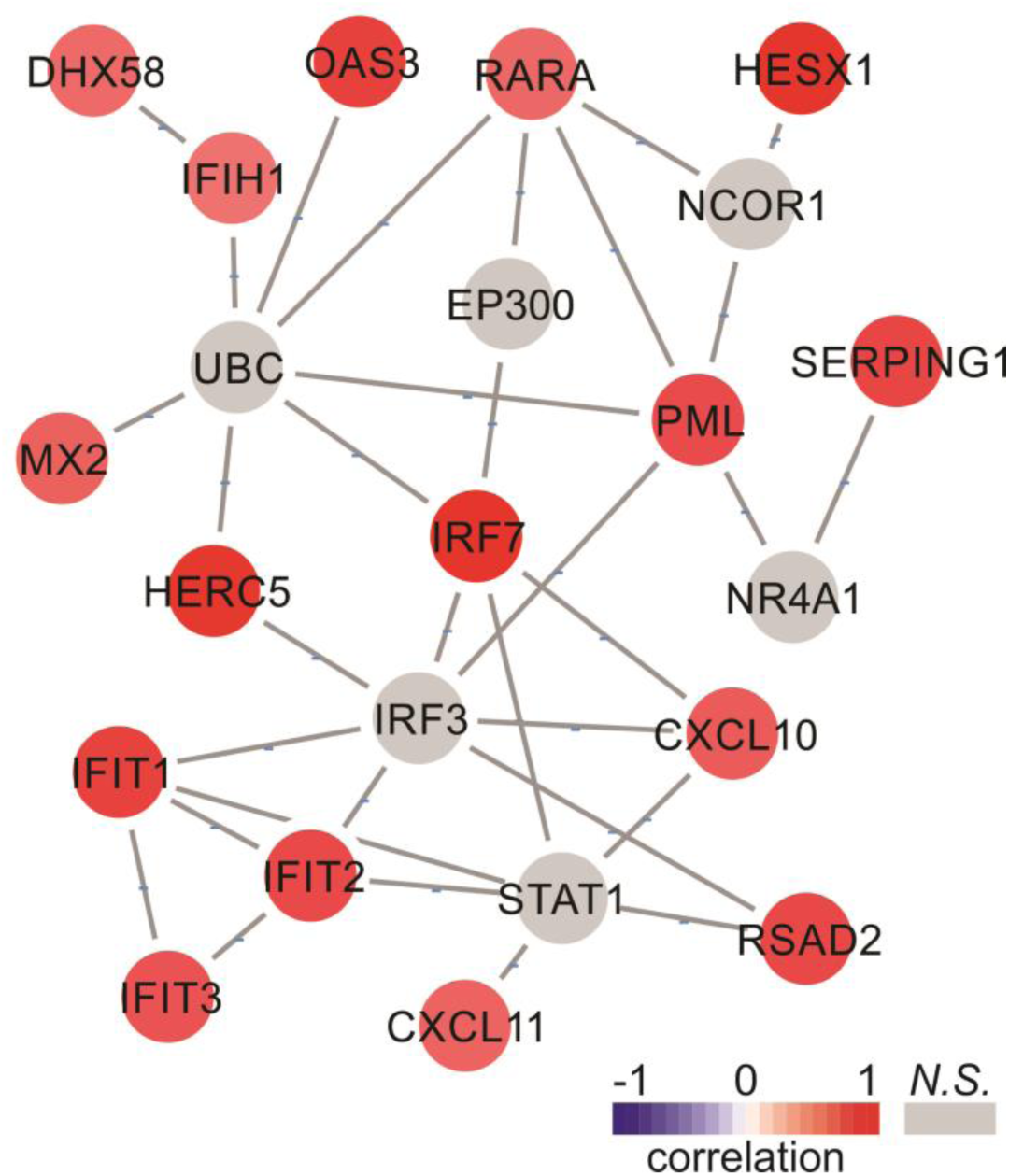
Network of genes whose expression is positively correlated with the amounts of CHIKV RNA. Minimum Networks constructed using the gene sets that presented a positive NES score with NetworkAnalyst tool. Red nodes represent genes that are positively correlated to CHIKV RNA and the color scheme indicates its strength. The gray nodes were added by NetworkAnalyst and are not part of the correlated genes.

The expression of several eukaryotic translation initiation factors (eIFs) was negatively correlated with the amount of CHIKV RNA (Fig. 3), indicating that these genes could play a major role in viral replication. The eIFs are important proteins that control translation initiation process, which is a major step for viral protein synthesis (20). A network with eIF genes was constructed (Fig. 3b) among which *EIF4B, EIF3L, EIF3E* and *EIF2AK2* presented the highest negative correlation with the amounts of CHIKV RNA (Fig. 3a). Only *EIF2AK2* gene also known as PKR had a positive correlation to CHIKV RNA in the blood of the infected individuals. It is well known that this protein could be activated by binding virus-derived double stranded RNA (dsRNA). This IFN-induced dsRNA-dependent serine/threonine-protein kinase inhibits viral replication by phosphorylating EIF2S1 (21) and shuts down cellular and viral protein synthesis.

**Figure 3.**
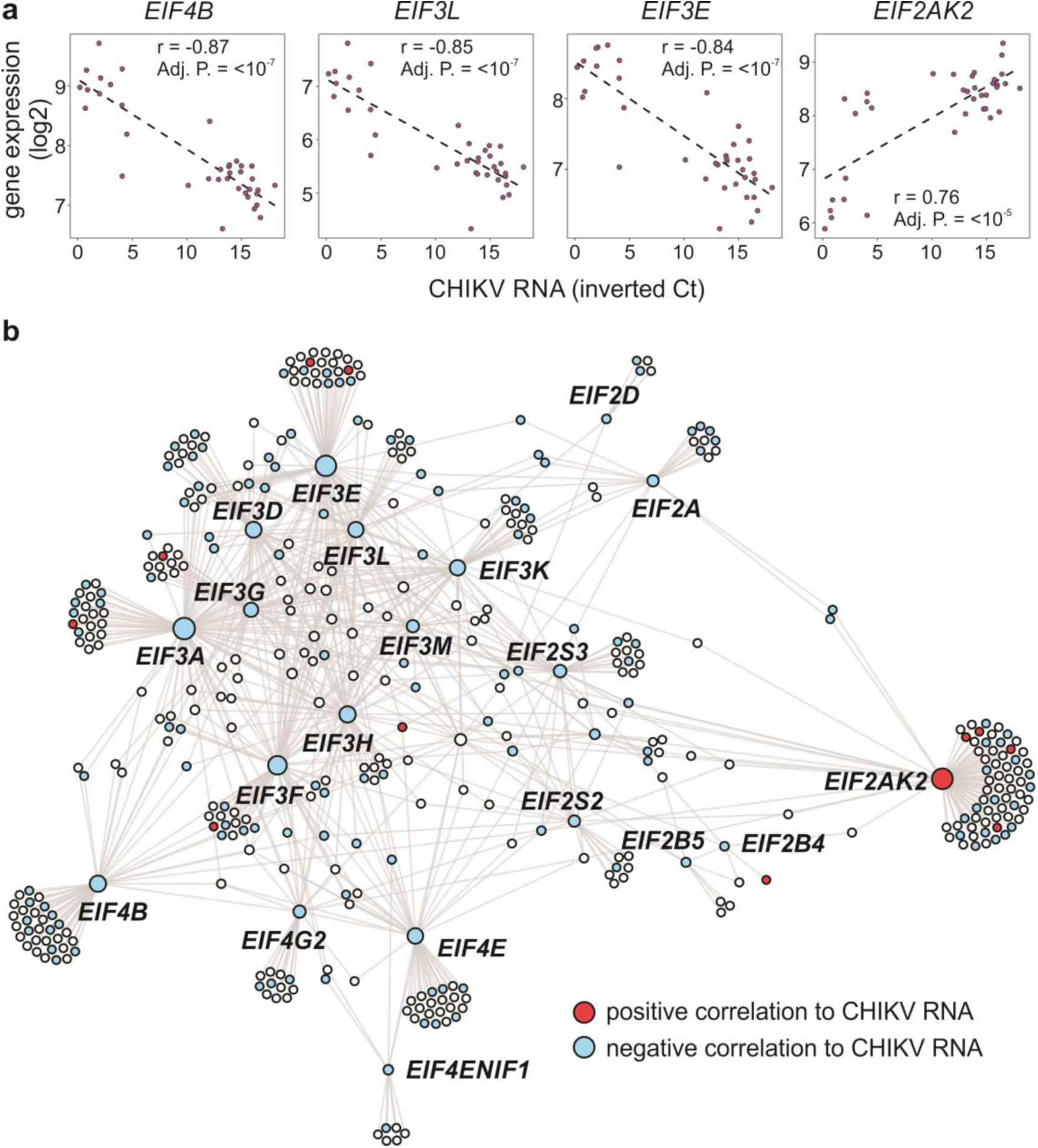
Eukaryotic Initiation Factor (eIF) genes association with CHIKV RNA. (a) Representation of the *eIF* genes correlation to CHIKV RNA (inverted Ct). (b) Network constructed with *eIF* genes correlated with CHIKV RNA and the genes associated with them integrated with PPIs. The NetworkAnalyst.ca tool (InnateDB PPIs; First Order) was used to construct the network and the Gephi program for visualization. The color of the nodes represents if the correlation between gene expression and CHIKV RNA is positive (red) or negative (blue).

### Consistent transcriptome changes upon CHIKV infection

Due to the natural heterogeneity in human cohorts, differential expression analysis was performed between each infected patient and the group of healthy controls. The patients whose samples were collected in the initial days post onset of symptoms presented higher numbers of DEGs (differentially expressed genes) detected by RNA-seq analysis and a higher mean of CHIKV RNA in their blood samples (Fig.4a). The DEG list of each patient was subsequently combined into a meta-volcano plot (Fig. 4b). This plot displays the number of genes whose expression was consistently altered in most patients. Approximately 70% of the CHIKV-infected patients presented down-regulation of 123 genes and up-regulation of 382 genes (Fig. 4b). *APOBEC3A, IFI44* (14), *OAS3* are the most represented up-regulated genes and *NT5E* (also known as CD73) *PTGS2, SNORD3C, EEF1A1P13* are the most represented down-regulated ones among the patients. APOBEC3 family members are important cytidine deaminases that control HIV replication (22) and have not been described till now in the CHIKV context. Since APOBEC3 genes are also considered ISGs, CHIKV infection are expected to induce this gene. Interestingly, the analysis also identified the *NT5E*/CD73 that is an ecto-5′-nucleotidase that has been described as an important molecule to recover the endothelial barrier during the dengue 2 infection leakage (23).

**Figure 4.**
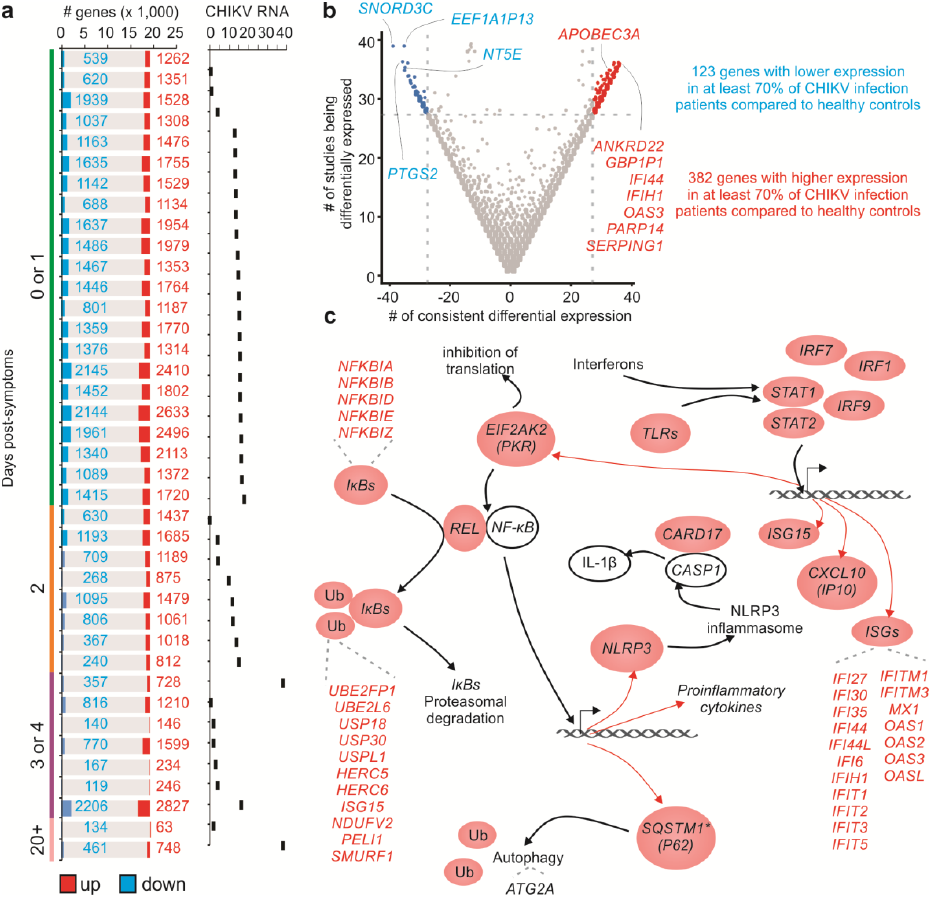
Differentially expressed genes among CHIKV samples. (a) Each infected patient’s profile was compared against healthy samples to exhibit the number of up- and down-regulated genes. The colors represent the number of up-regulated (red) or down-regulated (blue) genes and the respective number of genes in each group as indicated. The graph of bars to the right indicates the CHIKV RNA amounts in each patient separately. (b) The meta-volcano analysis showed genes that were consistently differentially expressed (log2 fold change > 1 and adjusted P value < 0.05) in at least 70% of the comparisons from infected samples versus healthy samples. The x-axis represents the number of consistent differential expressions of each gene while the y-axis represents the number of studies in which the genes are classified as a DEG. (c) Graphical representation of the putative relationship between some of the genes that were classified as consistently up-regulated in (b).

Through literature curation of the CHIKV gene signature, we created a network that provides novel insights into CHIKV immunobiology (Fig. 4c). CHIKV infection could induce the expression of several ISGs, as well as the activation of TLRs, *EIF2AK2* and IkBs. Up-regulation of *STAT1, STAT2, IRF9, IRF1* and *IRF7* can induce the expression of ISGs, which are important antiviral effectors. Interestingly, the CHIKV infection led to the up-regulation of several proinflammatory genes such as IκBs related genes, which are important to NFκB activation and proinflammatory cytokines production when phosphorylated. It is integral to mention here that we also detected the up-regulation of *REL*, which is a subunit of NFκB.

We also detected DEGs related to NLRP3 inflammasome. *Nlrp3* is transcriptionally regulated to guarantee high protein levels for activation of the NLRP3 inflammasome. Its activation could lead to autophagy (*SQSTM1*), apoptosis of infected cells and to other cytokines production (*CARD17*) in response to the infection (Fig. 4c). It has been shown recently that the activation of NLRP3 leads to IL-1β and IL-18 production and was linked to the alphaviral disease severity in an animal model (24).

The random forest method, a machine learning approach, was also employed to rank the importance of the 505 DEGs mentioned in the meta-volcano analysis (Fig. 4b) in predicting the CHIKV infection status. Fig. S3 displays the top 31 that better distinguish between infected patients and healthy control samples. The gene *SERPING1* presented an increased expression in HIV patients’ samples and could also be considered a CHIKV marker (25). Significantly, SERPING1 is a C1 complement protein inhibitor (26) and could be involved in signaling at the sites of inflammation contributing to tissue damage and disease severity as was suggested in the case of Ross River Virus infection. (27).

**Figure S3.**
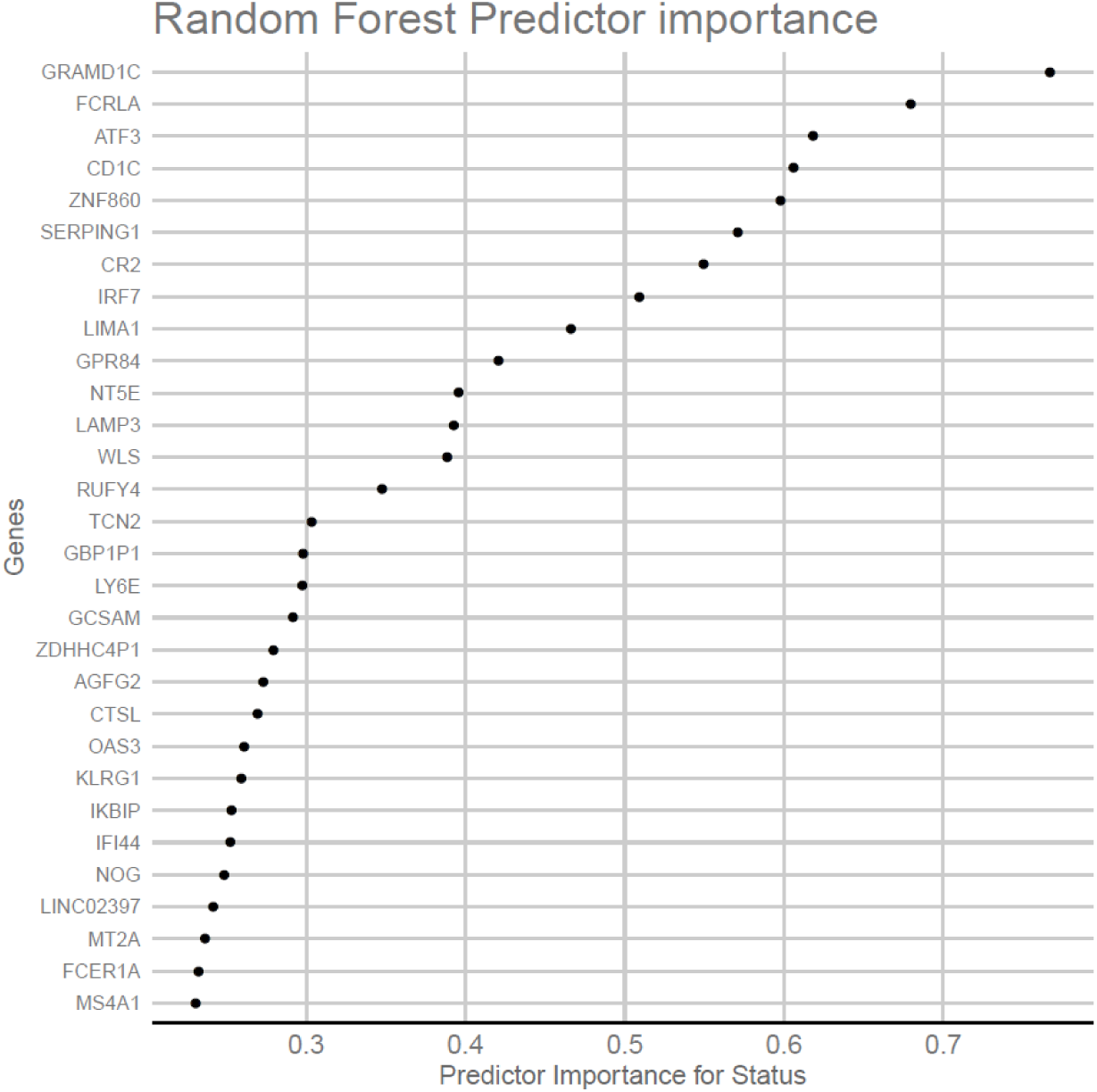
Random forest predictive analysis. The genes were sorted by the decreasing order of the Predictor Importance of Status used by random forest in order to prioritize the genes that most distinguish between infected and healthy samples.

We next performed single sample GSEA analysis using the log2 fold-change values of each patient compared to the group of healthy controls as ranks and the BTMs as gene sets (Fig. S4). Similar to the results presented in the Fig. 2, we observed that the up-regulated BTMs were related to innate immunity and anti-viral responses involving dendritic cells and monocytes activation.

**Figure S4.**
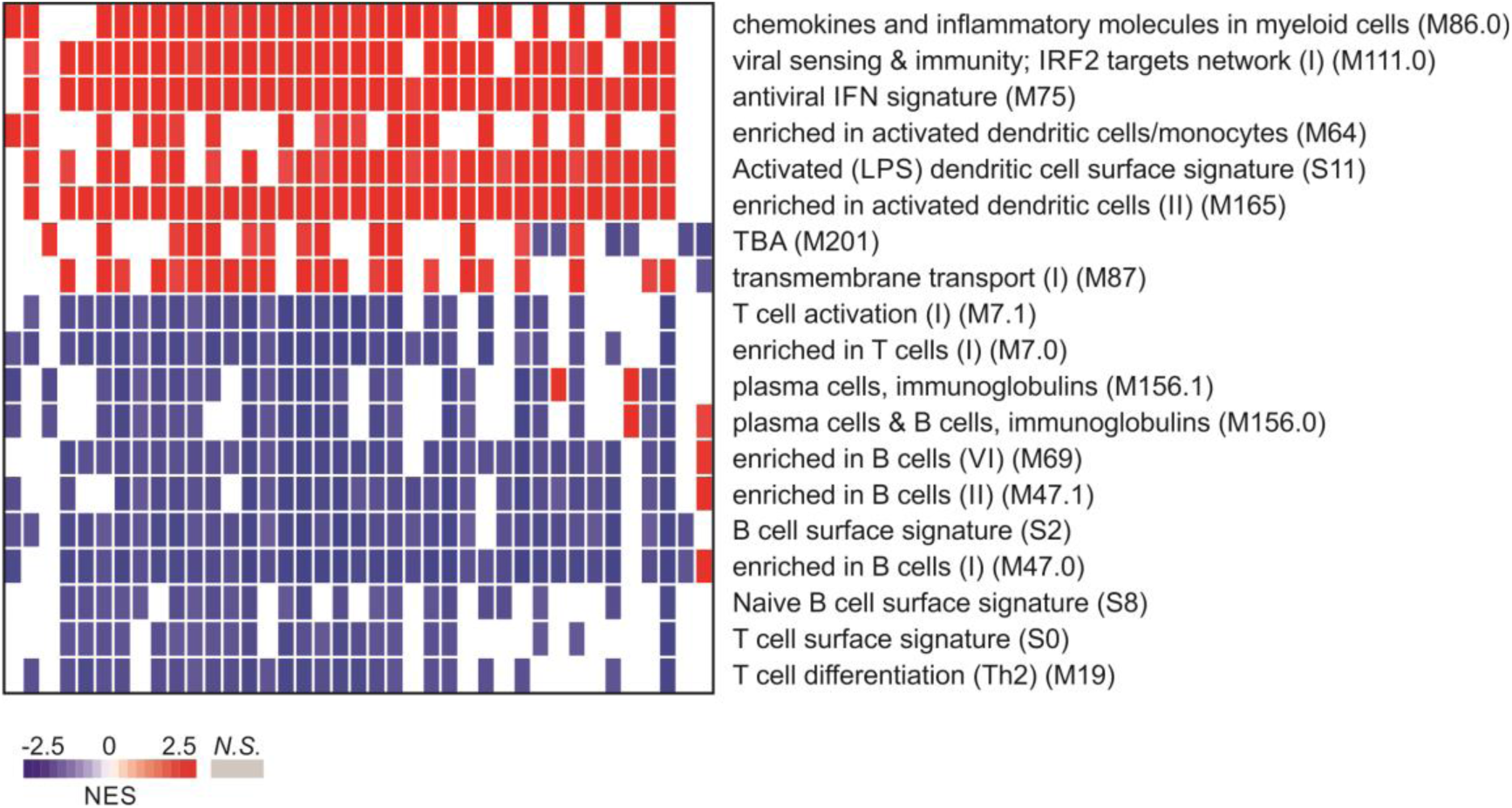
Gene Set Enrichment Analysis of BTM Pathways. GSEA was performed for each infected sample against the healthy control to retrieve a result of BTM pathways most consistently altered. The BTMs were used as gene sets and their respective log2 fold-change results as rank. Each column represents the results from GSEA comparison between each infected sample against the healthy control. The pathway names are shown in the right side of the heatmap. Pathways profiles were ordered by the mean of the NES score across all patients.

### Potential Signatures of CHIKV-induced Arthralgia Chronicity

We split the patients (n = 13) who agreed to return for clinical follow-up examinations into those with chronic and non-chronic arthralgia (see methods). These two groups possess no difference in the amounts of CHIKV RNA in the serum (Fig. S5a). Compared to the group of healthy controls, we found that a total of 1,262 and 1,862 genes were consistently differentially expressed in most of the chronic and non-chronic patients respectively (Fig. S5b). Of those, 514 were commonly up-regulated in both groups whereas 337 genes were down-regulated in both groups (Fig. S5c). Additionally, there is a high positive correlation between the mean log2 fold change of chronic and non-chronic patients when compared to healthy controls (Fig. S5d). However, few genes presented an expression with inverted behavior between chronic and non-chronic patients (Fig. S5d). *HLA-DRB5* that could be related to antigen peptides presentation is one example of genes that was up-regulated in most non-chronic samples and down-regulated in most chronic samples (Fig. S5d). Genes such as *DDX3Y, EIF1AY*, and *LINC00278* were up-regulated in most chronic samples and down-regulated in most non-chronic samples. The *EIF1AY* gene belongs to the family of eukaryotic translation initiation factors (and showed in Fig. 3). Interestingly, the *DDX3Y* is an RNA helicase that was described as an important effector of the herpes virus replication and propagation (28) but had not been described in the context of CHIKV.

**Figure S5.**
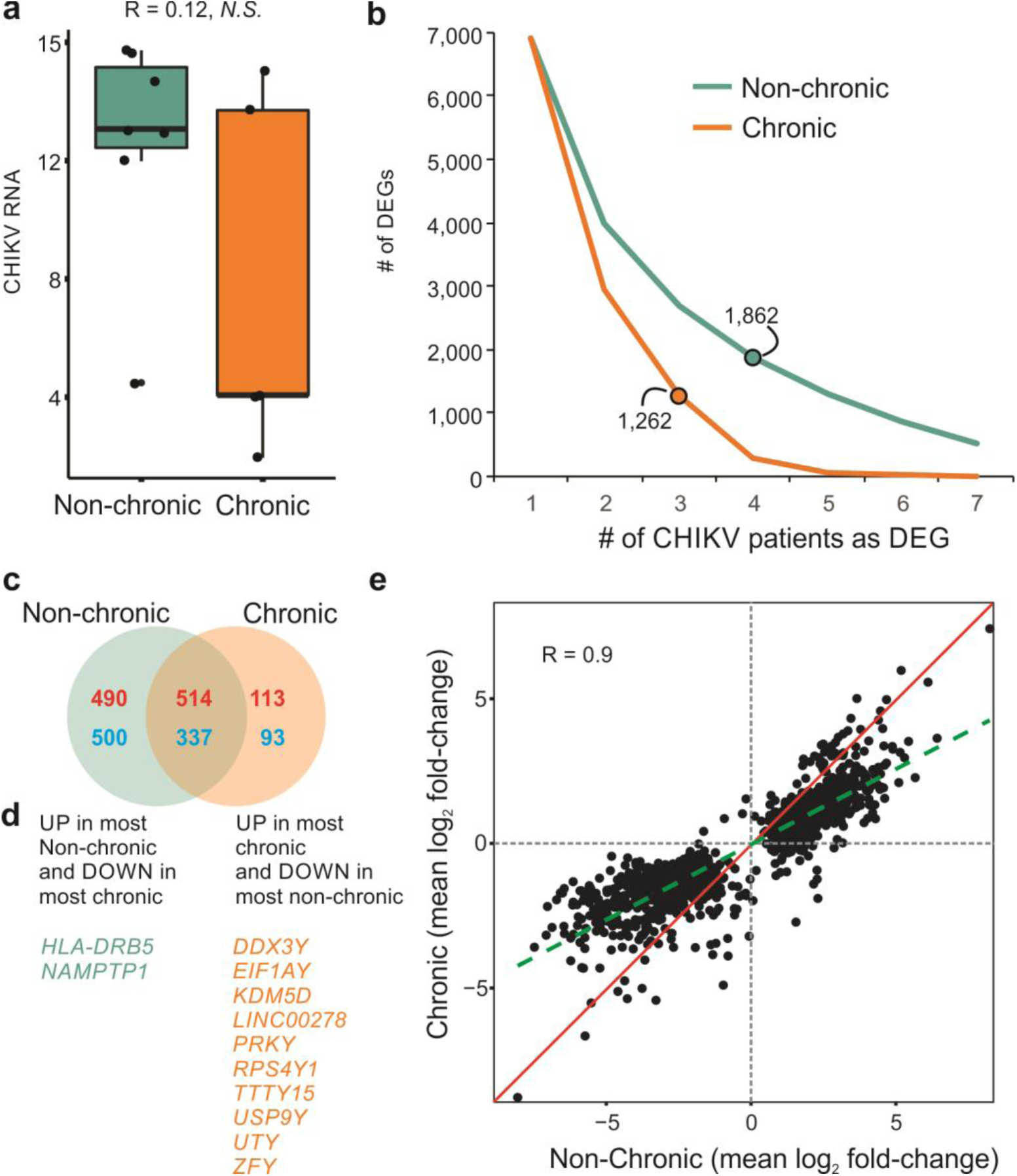
Transcriptome changes in chronic and/or non-chronic subjects compared to healthy individuals. (a) Comparison of the amounts of CHIKV RNA (inverted Ct) between chronic (n=6) and non-chronic (n=7) subjects. (b) Number of genes consistently differentially expressed in chronic (brown) and non-chronic (green) subjects. The x-axis represents the number of CHIKV-infected subjects used in each analysis and the y-axis indicates towards the number of DEGs consistently found. The dot marked in the plot represents the number of DEGs in at least 70% of the samples in each group. (c) Representation of differentially expressed genes (DEGs) in at least 70% of the chronic and/or non-chronic patients compared to healthy subjects. (d) Genes that overlapped in (c) but present an inversed behavior depending on the group.

### Modular expression analysis of CHIKV infection

We performed a gene co-expression network analysis using the expression profiles of all the patients and healthy controls. CEMiTool (29) identified eight co-expression modules containing 74 to over 2,000 genes (Fig. 5a). The expression activity of some of these modules were altered in healthy controls or in patients with different days after the symptoms onset (Fig. 5b). The module M8, which shows higher expression in patients with two to four days of symptoms was enriched for monocytes and neutrophils (Fig. 5c). These immune cell subsets are known for being related to the disease (30). Moreover, CEMiTool can integrate the gene modules with protein-protein interaction data and identify important hubs within each module. In module M8, we found as one of the hubs the gene *DDX58* (also known as *RIG-1*) that has already been linked to CHIKV disease (30) and as a virus replication inhibitor in an animal model for CHIKV infection (31).

**Figure 5.**
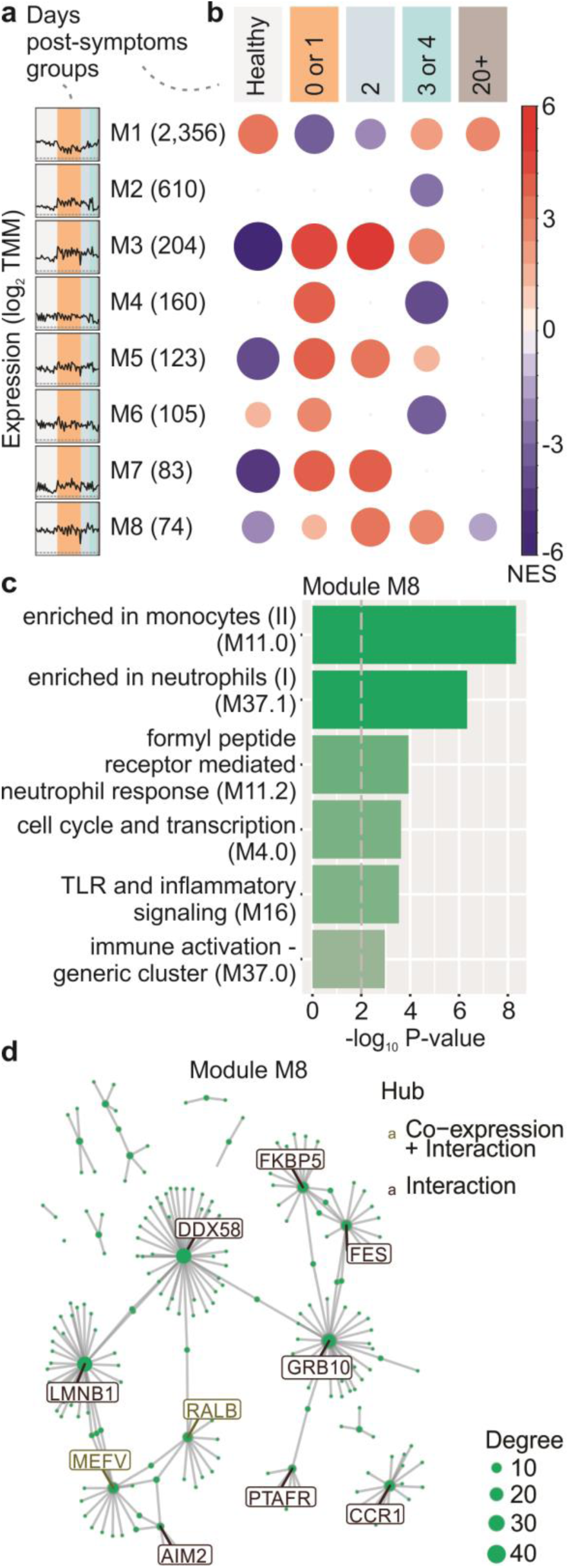
Modular analysis of CHIKV infection. (a) Expression profile of all the eight modules returned by CEMiTool. The colors represent the days post onset of symptoms and the black line represents the mean expression of all the genes inside the module. (b) Gene set enrichment analysis showing the module activity for each subgroup. The size and color of the circle represents the normalized enrichment score (NES). (c) Over representation analysis of module M8 using BTM pathways as gene sets. The pathways were ordered by significance as indicated in the x-axis. (d) Gene network of module M8 for the top ten most connected genes (hubs) represented as nodes and its protein-protein and co-expression interactions as edges. The size of the node represents its degree of connectivity.

### Comparing CHIKV signature with Rheumatoid Arthritis and Dengue infection signatures

To check the extent to which the gene signature was specific to CHIKV infection, the CHIKV signature was compared with those from another viral infection (dengue virus) or from an auto-immune related inflammatory disease (rheumatoid arthritis or RA). We re-analyzed two publicly available blood transcriptome datasets and identified the genes whose expression was altered in the dengue infected patients and RA patients compared to healthy controls. A total of 949, 632 and 302 genes were identified as being up-regulated only in RA, dengue infection and CHIKV infection respectively (Fig. 6a). Among those, seven up-regulated genes were shared by all the three signatures, including the following: *OAS1, C1QB, ANKRD22, IRF7, CXCL10, IFI6* and *IFIT3* (Fig. 6a). More than 300 genes were exclusively up-regulated upon CHIKV infection, including the inflammasome-related *NLRP3* genes related to NFκB and genes that belong to the Th2 response (*ILI31RA, IL4I1*) (32) (Fig.6a).

**Figure 6.**
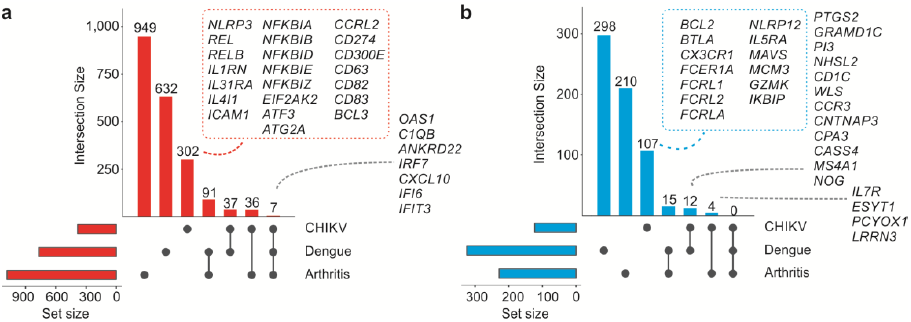
Comparing CHIKV signatures with those from rheumatoid arthritis and dengue infection. (a) Genes up-regulated in patients infected with either CHIKV or dengue or rheumatoid arthritis (RA) patients compared to their respective controls. The size of the bar represents the quantity of up-regulated genes detected in one or more groups (indicated as a gray circle in the graph below). Representative up-regulated genes are displayed. The gray circles below the bars indicate the groups that share the same up-regulated genes. (b) The same approach as in (a) but for the down-regulated genes.

Although no gene was down-regulated in all the three conditions, genes such as *IL7R, ESYT1, PCYOX1* and *LRRN3* were commonly down-regulated in CHIKV infection and RA samples (Fig. 6b). The cytokine IL-7 is crucial for the survival of naïve and memory T cells, which are important effectors against several pathogens including viral infections. The receptor IL-7R is expressed on the surface of these cells and was shown in an animal model to prevent the chronicity induced by lymphocytic choriomeningitis virus due to the enhancement of CD8 T cells’ response and prevention of its exhaustion (33). We identified a specific CHIKV signature composed of 107 down-regulated genes that includes *CX3CR1, BTLA, BCL2, GZMK* and three genes that encode Fc receptor-like glycoproteins (Fig. 6b) that could be related to the lymphopenia induced by CHIKV infection (34). The down-regulation of CX3C chemokine receptor 1 (*CX3CR1*) seems to agree with the results of this study that showed that several T cell effector/differentiation genes were down-regulated in CHIKV infection. CX3CR1 has been considered as an important marker of CD8 memory cells (35). Moreover, *BCL-2* down-regulation could be a mechanism of CHIKV host response to induce apoptosis of infected cells.

As the inflammasome-related genes are exclusively up-regulated in CHIKV infection and was described before in this context (24), we decided to provide a deeper proof-of-concept related to these findings. The murine bone marrow-derived macrophages were infected with CHIKV virus and the readouts of the inflammasome activation were measured. We used a specific dye that binds to active caspase-1 (FAM-YVAD) to assess caspase-1 activation and it was observed that CHIKV induce caspase-1 activation as measured by the percentage of FAM-YVAD+ cells and the integrated mean of fluorescence (iMFI) (Fig. 7a). Caspase-1 activation was further assessed by western blot and detected caspase-1 cleavage as shown by the presence of Casp1 p20 in cells infected with MOI of 5 (Fig. 6c). We found that CHIKV infection at MOI of 1 and 5 induces IL-1β production and LDH release (Fig. 6d). IL-1β production was dependent on the inflammasome because Casp1/11 deficient macrophages fail to trigger IL-1β production in response to the infection (Fig. 6e). These data in conjunction indicate that the inflammasome is activated in response to the infection, and thereby support the robustness of our transcriptional analyses.

**Figure 7.**
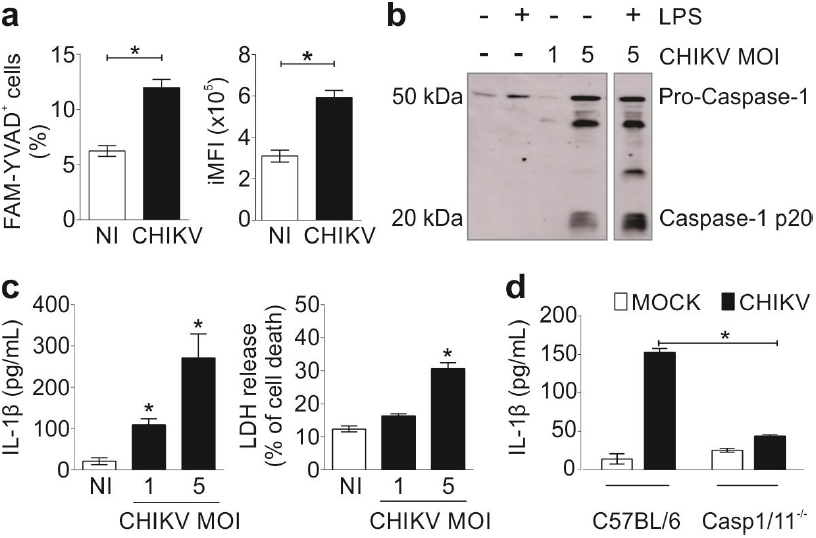
CHIKV activates inflammasome in murine macrophages. (a) Mouse bone marrow-derived macrophages (BMDMs) were infected with CHIKV with a MOI of 5 or not infected (NI), and after 24 hours of infection the cells were stained for active caspase-1 with FAM-YVAD. The percentage (left) and integrated mean of fluorescence (iMFI) of activated cells (right) is demonstrated. (b) After 24 hours of infection, supernatants were harvested from non-infected (NI) or CHIKV-infected BMDMs, treated with LPS as depicted and levels of cleaved caspase-1 (p20) were detected by western blotting in the supernatant. (c) LPS-primed BMDMs were infected CHIKV with a MOI of one or five. After 24 hours of infection, the levels of IL-1β (left) and LDH (right) in the cell-free supernatants were measured. (d) Production of IL-1β 24 hours post-CHIKV infection by BMDMs from wild type (C57BL/6) or Casp1/11 deficient mice. Data from (a), (c) and (d) are represented as the mean ± SD of triplicate samples and are representative of the three independent experiments that yielded similar results. Statistical analysis was performed by student’s t-test. Asterisks indicate statistically significant differences. *P < 0.05.

## Discussion

CHIKV re-emergence in the past 15 years has been producing major epidemic outbreaks in Asia, Africa, the Indian Ocean and more recently in the Americas after decades of intermittent outbreaks (36). Despite considerable progress in understanding the infection, much of the host–pathogen interplay remains obscure.

Systems biological approaches can provide comprehensive and unbiased dissections of the complex interactions between genes and proteins during an infection (15). Relevant studies related to the natural infection by CHIKV were recently published. However, these were limited to the analysis of a limited number of proteins (37) or on children infected with the virus (38). Since it is extremely difficult to acquire sufficient data from naturally infected individuals for use in systems biological models of analysis, this kind of study is relatively new. To our knowledge, we are the first to investigate the early host response to acute CHIKV infection in adults using a systems biological approach.

The most clinically relevant symptom related to CHIKV infection is peripheral symmetrical joint pain (primarily inflammatory polyarthralgia). Manifested in several CHIKV cases, it can produce a high economic impact and severely affect the patients’ quality of life (11). The pain starts in the acute phase and can persist for years in up to 50% of the infected adult individuals (39). Notably, it was proposed that CHIKV infection produced autoimmune sequelae (40). Although polyarthralgia is rarely detected (5–11%) in children, other kinds of severe manifestations can occur (41). Since there are neither effective treatments nor licensed vaccines against CHIKV infection, understanding the molecular mechanisms of this complex infection is essential for the development of efficient therapies and even vaccines.

Consistent with other reports, it was observed the up-regulation of several genes that play roles in the antiviral immune response, and many of them are ISGs (14, 19, 42). Similarly, other reports that assessed the changes in immune cell subsets during CHIKV infection (38, 43) support the findings of this study regarding the most abundant up-regulated genes being related to neutrophils and myeloid populations and especially dendritic cells and monocytes. The monocytes have been reported recently as important inflammatory and regulatory mediators of the innate immune response to different arboviruses (15, 44). In corroboration with these findings, we also showed the exclusive up-regulation of inflammasome-related genes. Additionally, the induction of inflammasome activation in macrophages infected with CHIKV *in vitro* was observed. Importantly, these data account to explain our previous demonstration that NLRP3 inhibitors can interfere with the pathogenesis of CHIKV virus (24).

Moreover, the negative and strong correlation between CHIKV RNA amounts and most eIF family member genes observed here have not been described previously in CHIKV infections. Furthermore, it is speculated whether the down-regulation of other eIF family members observed in these results in response to the infection could be an important host defense mechanism against the virus replication. We consider that the eukaryotic translation initiation factors are attractive markers of the CHIKV arthralgia chronicity and could be better exploited as novel broad-spectrum antiviral targets.

The CHIKV related arthralgia symptom is very similar to the rheumatoid arthritis (RA) but there are some important clinical and immunological characteristics that differ between these diseases and the different molecular signatures between both CHIKV and RA can be shown here. Previous data showed significant concordance between rheumatoid arthritis gene signatures and a mouse model of CHIKV infection/arthritis (13). Here, we are presenting a more detailed comparison between both diseases in humans.

On the other hand, when compared to the differential gene expression of DENV infected patients, the data in this study shows specific molecular gene signatures in response to CHIKV. It is interesting to note in this context that although previous infections with ZIKV and DENV were detected in the group of non-CHIKV-infected volunteers, specific responses for CHIKV infection were still possible to identify.

Moreover, the comprehensive molecular data in this study showed novel molecules that could play important roles in the acute infection of CHIKV infection thereby providing new candidates as targets for therapies against this incapacitating disease.

## Methods

### Sample collection and clinical information

Blood samples were collected from subjects reporting arbovirus-like symptoms in the Brazilian states Sergipe (n=39 infected and n=15 controls) and São Paulo (n=5 controls). The study was approved by the Ethics Committees from both the Department of Microbiology of the Institute of Biomedical Sciences of the University of São Paulo and the Federal University of Sergipe (Protocols: 54937216.5.0000.5467 and 54835916.2.0000.5546, respectively). Clinical and socio-demographic data were collected through a questionnaire filled by the patients. Total peripheral blood was collected in Tempus tubes or in heparin treated tubes which were then transferred to tubes containing RNALater (Thermo Fisher) and stored at −80°C. A subset of the patients (n=13) agreed to return for a clinical follow up. These CHIKV infected patients were under treatment and their joint manifestations were monitored quarterly. Patients that presented joint pain associated or not with joint or periarticular edema of onset or significant worsening (in cases of previous joint disease) after three months of acute febrile syndrome were considered chronic patients.

### Molecular Diagnostics

Real-time RT-PCR was performed to test for CHIKV, ZIKV and DENV as previously described (17, 45). Nucleic acid extraction was performed using the QIAamp Viral RNA Mini Kit (Qiagen, Valencia, CA, USA) and carried out according to the manufacturer’s instructions. Molecular detection of DENV, CHIKV, and ZIKV was performed by using the SuperScript III Platinium One-Step qRT-PCR kit (Invitrogen). We performed the samples’ amplification of the RNAse P in parallel to evaluate the extracted RNA quality. RNA Real Time RT-PCR reactions consisted of a step of reverse transcription at 50 °C for 15 min of the enzyme activation at 95 °C for 2 min, and 45 cycles at 95 °C for 15 s and 60 °C for 1 min for hybridization and extension with the use of ABI 7500 equipment (Applied Biosystems). A cut-off value of Ct (Cycle Threshold) 37 was chosen as the CDC reference assay.

### Serological Diagnostics

Serum samples were evaluated with a commercial semi-quantitative ELISA kit (enzyme-linked immunosorbent assay), which detects anti-CHIKV IgM and anti-CHIKV, Zika and Dengue IgG antibodies. All procedures were carried out according to the manufacturer’s instructions (Euroimmun, Lubeck, Germany). Briefly, sera were diluted in sample buffer and incubated at 37 °C for 60 min in a microplate well together with a calibrator, and positive and negative controls provided by the manufacturer. The optical density (O.D.) was measured in an Epoch Microplate Spectrophotometer (BioTek, Vermont, USA) and the results were calculated according to the manufacturer’s instructions. Samples with ratio values (Extinction of the control or patient sample / Extinction of calibrator) below 0.8 and above 1.1 were considered negative and positive samples, respectively. Samples with ratio values between 0.8 and 1.1 were considered inconclusive.

### RNA-Seq experiments

Total RNA from Tempus tube samples were purified with the Tempus™ Spin RNA Isolation Kit (Invitrogen). Total RNA from samples collected in heparin vacutainer tube and maintained in RNAlater were purified using the Ribo Pure™ blood isolation kit (Invitrogen). Removal of rRNA and globin-encoding mRNA, RNA fragmentation, cDNA generation, adapter ligation and PCR amplification were performed using the TruSeq stranded total RNA with ribo-zero globin sample preparation kit (Illumina). The sequencing of transcriptome libraries was performed on the Illumina HiSeq 1500 platform (Illumina, San Diego, CA). Libraries were prepared using TruSeq™ Stranded RNA Sample Preparation kit with Poli(A)+ selection, quantified through qPCR and sequenced using HiSeq SBS V4 kit (2 x 125 bp paired-end reads).

### RNA-Seq data analyses

Raw paired-end reads were preprocessed for quality control. The Trimmomatic software, version 0.36 (46) has been used to remove adapters, to trim the 5′ and 3′ ends with mean quality score below 25 (Phred+33), and to discard reads shorter than 35 bp after trimming. Paired-end reads mapping to PhiX Illumina spike-in were removed using Bowtie 2, version 2.2.5 (47). After preprocessing, the high quality paired-reads were mapped into the reference genome *Homo sapiens*, version GRCh38.p10 build 38 with the TopHat2 program (48), based on 57,685 genes from *Homo sapiens*, excluding the rRNA and globin genes. For the TopHat2 alignment, we considered the following parameters: minimum intron size (30pb), number of mismatches per read (3pb), number of gaps per read (3pb), --very-sensitive, maximum insertion size deletion (3bp), maximum paired-reads distance (200pb) and only concordant uniquely mapped reads (approximately of 92% mapped reads) were used in further analyses. To quantify the gene abundance of mapped paired-end reads in each sample, we used the featureCounts tool from Bioconductor Rsubread package (49). The total number of read counts per gene was obtained from the RNA-seq expression. Normalization of the gene counts was performed with counts per million normalization (CPM), which accounts for differences in library size and adjusts for GC content and gene length. Only genes with counts per million (CPM > 1) in at least two sample replicates were used in the analyses. To estimate the source of variability of experimental effects, an unsupervised analysis was performed with the R package “PVCA” (50).

The identification of differentially expressed genes (DEGs) were carried out using Bioconductor package edgeR. DEGs were identified following two criteria: (i) adjusted p-value < 0.05 and (ii) fold change > 2. DEGs shared among CHIKV patients were found using the MetaVolcano webtool (https://metavolcano.sysbio.tools/).

The Pearson correlation test (|R| > 0.5 and adjust p-value < 0.01) was used to identify associations between genes and inverted Ct. R package “randomForest” was used to perform a Random Forest analysis to classify the importance of genes correlated to CHIKV RNA in predicting CHIKV infection.

Regarding the pathways that may be related to the progression of the disease, a Gene Set Enrichment Analysis (GSEA) was performed using as ranks the correlation between the genes and inverted Ct. A set of Blood Transcriptional Modules (BTM), previously identified by our group (51) through large-scale network integration of publicly available human blood transcriptome, was used as gene sets.

### Gene Co-expression and Network Analysis

We performed the gene co-expression analysis using the R package CEMiTool (52). For this analysis, we normalized the expression data using TMM (Trimmed Mean of M-values) and transformed it to log2 scale. We followed the default parameters with a variance filter of 0.2.

To gain a systems-level understanding of the patterns of a certain disease, one of the steps required is the construction and analysis of the network involving the most interesting genes. For this, we used NetworkAnalyst (https://www.networkanalyst.ca/) with the protein-protein interaction (PPI) database based on InnateDB. To improve the visualization, the software Cytoscape (https://cytoscape.org/) and Gephi (https://gephi.org/) were also used.

### Meta-analysis of Dengue and Rheumatoid Arthritis Transcriptome studies

The transcriptome datasets of patients with either rheumatoid arthritis (RA) or Dengue infection were downloaded from the Gene Expression Omnibus (GEO) under the accession GSE51808 and GSE51808. DEGs between RA patients and healthy controls and between Dengue-infected patients and non-infected subjects were identified using the limma package (Adjusted P-value < 0.05 and fold-change > 1.25).

### Bone marrow–derived macrophage preparation and infections

Bone marrow–derived macrophages (BMDMs) were prepared using tibia and femur from 6- to 12-week-old mice as previously described (53). Wild type (WT) C57BL/6 mice (Jackson Laboratory, stock number 000664) and Casp1/11^-/-^ (54) derived from C57BL/6 mouse strains were used. All mice were bred and maintained under specific-pathogen-free conditions at the Animal Facilities of the Medical School Ribeirão Preto (FMRP-USP). Mice experiments were conducted according to the guidelines of the institutional committee for animal care (Protocol number 14/2016).

The virus strain CHIKV BzH1 was used and virus stocks were produced by infecting Vero cells with a MOI of 0.1. Conditioned media used for mock infections was prepared from uninfected Vero cells in a similar manner. The MOI of 5 and the time point of 24 hours of infection was used in most of the *in vitro* experiments using BMDMs, unless otherwise stated in the figure legends.

### Quantification of IL-1β secretion and cell death assay

For *in vitro* cytokine determination and LDH assay, BMDMs were seeded at a density of 2 × 10^5^ cells/well in 48-well plates and pre-stimulated with 500 ng ml^-1^ of LPS (tlrl-peklps; InvivoGen) for 3 h, and subsequently infected with CHIKV. The cytokines in the supernatants were measured using mouse IL-1β ELISA kit (BD Biosciences) according to the manufacturer’s instructions. LDH measurement was performed with CytoTox 96 Non-Radioactive Cytotoxicity Assay (Promega) following manufacturer’s instructions. The positive control for complete cell lysis and normalization was 9% Triton X-100 (Fisher Scientific), it was incubated with cells for 15 min. The percentage of LDH release was calculated as (mean OD value of sample / mean OD value of Triton X-100 control sample) × 100 and is shown in the figures as the percentage of cell death compared to TritonX-100 (%).

### Caspase-1 evaluation by western blot analysis and endogenous caspase-1 staining using FAM-YVAD–FMK

In order to measure active caspase-1 we used FLICA assay. Briefly, 10^6^ BMDMs were seeded in 12-well plates overnight, and then infected with CHIKV at a MOI of 5 for 24 hours. As a positive control, we used 20 μM of nigericin (Sigma-Aldrich) for 40-60 minutes. After that, cells were harvested and stained for 1 h with a green fluorescent dye that binds specifically to active caspase-1, FAM-YVAD-FMK (Immunochemistry Technologies), following the manufacturer’s instructions. The data were acquired on a FACS ACCURI C6 flow cytometer (BD Biosciences) and analyzed with the FlowJo software (Tree Star).

For the western blot 10^6^ BMDMs were seeded in 6-well plates overnight, and then primed with 500 ng ml-1 LPS (InvivoGen, tlrl-peklps) for 3 hours prior to infection with CHIKV. The supernatants were collected, and precleared supernatants were concentrated 3 times in speedvac. They were boiled in Laemmli buffer, resolved by SDS-PAGE and transferred (Semidry Transfer Cell, Bio-Rad) to a 0.22-μm nitrocellulose membrane (GE Healthcare). The membranes were blocked in Tris-buffered saline (TBS) with 0.01% Tween-20 and 5% nonfat dry milk. The rat monoclonal antibody to caspase-1 p20 (1:250, Genentech, 4B4) and specific horseradish peroxidase–conjugated antibodies (1:3,000, KPL, 14-16-06 and 14-13-06) were diluted in blocking buffer for the incubations.

Data were plotted and analyzed with GraphPad Prism 6.0 software (GraphPad, San Diego, California). Multiple groups were compared by two-way analysis of variance (ANOVA) followed by the Bonferroni’s post-test were used. The differences in values obtained for two different groups were determined using an unpaired, two-tailed Student’s t test with a 95% confidence interval. Differences were statistically significant when the p value was less than 0.05.

## Acknowledgements

H.I.N. is supported by the São Paulo Research Foundation (FAPESP; grants 2017/50137-3, 2012/19278-6, and 2013/08216-2). A.S.S. is supported by Butantan Foundation, CNPq (Grant 443371/2016-4) and Brazilian Health Ministry. R.A. is supported by FINEP Grant 0116005600. D.R.R. has a postdoctoral fellowship from CNPq. J.C.A. has a postdoctoral fellowship from CAPES - Finance Code 001. M.P.C. has a PhD fellowship from FAPESP - 2016/08204-2. I.J.A is supported by São Paulo Research Foundation (FAPESP; grant: CEPID 2013/07467-1). P.L.H is supported by Butantan Foundation, CNPq 306992/2014-0 and Fapesp 2015/25055-8.

